# Deleterious imprinting of perinatal exposure to nano-polystyrene on inflammatory bowel diseases

**DOI:** 10.1101/2024.12.20.629591

**Authors:** Melvin Airaud, Amandine Roldan, Jeanne Le Cléac’h, Lisa Isoard, Emmanuel Mas, Marie Carrière, Sandrine Ménard, Frédérick Barreau

## Abstract

The tremendous and exponential production of plastics as well as its poor recycling leads to massive release in the environment questioning their outcomes on human health. Plastics are degraded into micro-plastic (MPL) and nano-plastic (NPL) that undergo weathering. MPL and NPL enter the human food chain via contaminated food, when animals are bred or vegetables grew on in plastic polluted environment and through the degradation of plastics food packaging. Ingested MPL or NPL can pass through epithelial barrier. This study focuses on consequences of oral perinatal exposure to polystyrene 50nm pristine or weathered (w) on intestinal disorders onset in offspring.

Mouse were forced fed daily with 1.25mg of PS50 or PS50w starting at 15 days of gestation until pups weaned (Post Natal Day PND21). Oral tolerance protocol was induced at PND15 (nutritional switch) with ovalbumin (OVA) whereas DSS-induced colitis was performed in PND63 offspring.

PS50 nor PS50w perinatal exposure did impair tolerance or immunization to OVA at PND15 in male or females. However, PS50 and PS50w perinatal exposure significantly worsened macroscopic score of DSS-induced colitis in both sexes. Perinatal exposure to PS50w appeared to be even more deleterious than PS50 for colitis exacerbation in males.

In conclusion, early life exposure to PS50 or PS50w has long-lasting consequences on gut physiology by increasing the colonic severity of colitis while it does not modify oral tolerance establishment in early life. Those preliminary results questioned the kinetic of perinatal PS50 and PS50w exposure outcome over the life course and introduce the notion of imprinting.

**Highlights:**

- Perinatal exposure to PS50 or PS50w does not affect oral tolerance establishment nor immune response to immunisation at PND15
- Perinatal exposure to PS50 or PS50w have long lasting consequences on intestinal inflammation and colitis. PS50w in male has even more deleterious outcomes.
- Restricted exposure to PS50 or PS50w during perinatal has a deleterious imprinting on adult offspring intestinal health

**Graphical abstract:** 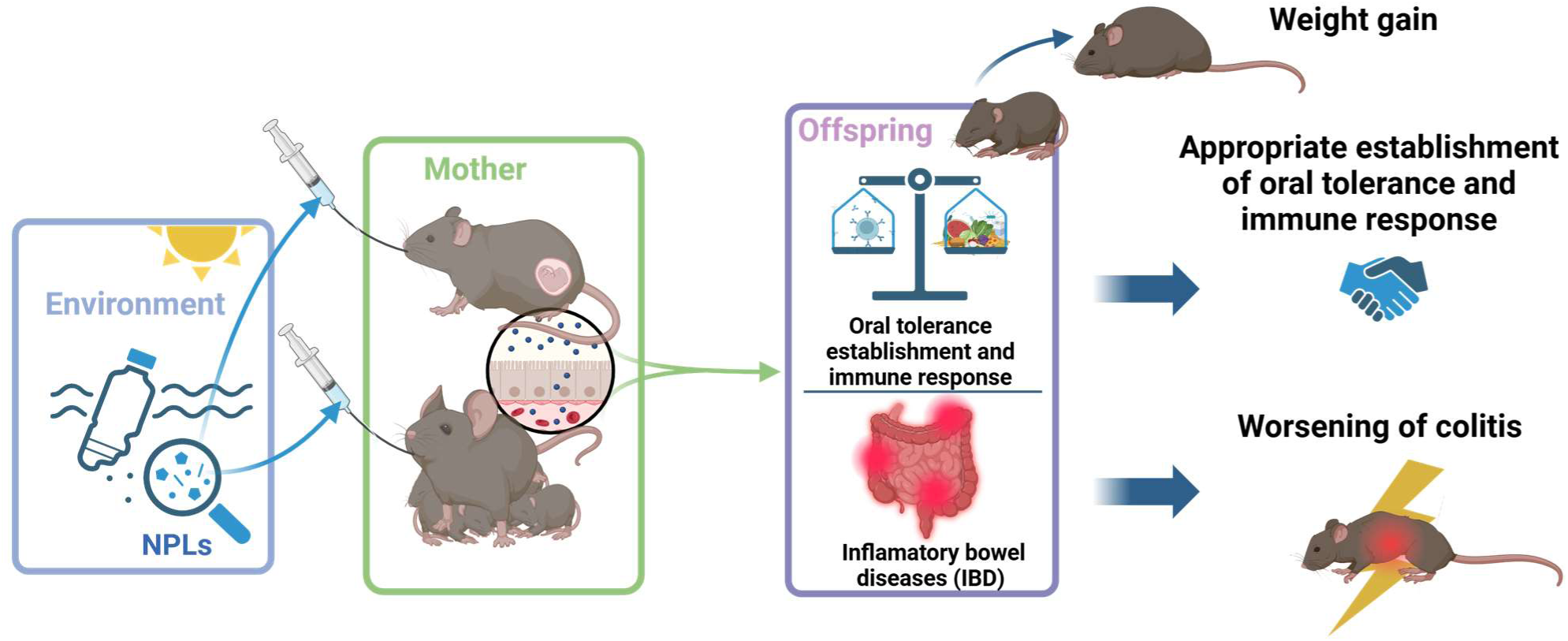

**Environmental implication:** The use of plastics in daily products leads to ubiquitous contamination of the environment and subsequently of food chains. Plastics persist for long periods in the environment where, submitted to climate, they fragment into nanoplastic particles (NPL). Indeed, these NPL are weathered due to their exposure to UV light, thermal, chemical, physical and biological stresses. Thus, human populations are exposed to pristine (NPLp) and weathered NPL (NPLw) in their daily lives, with unknown effect on intestinal health, including the diseases development like adverse Food Reactions (AFR) or Inflammatory Bowel Diseases (IBD). Given that these pathologies could originate in perinatal development and that pregnant and breastfeeding women are exposed to NPL on a daily basis, this study explores the impact of perinatally exposure to pristine or weathered polystyrene nanoplastics on the gut development and susceptibility.

## Introduction

Today, plastic represents one of the main environmental pollutant, due to the constant increase in its production over the last century. According to Organization for Economic Co- operation and Development report, from 0.5 million metric tons (Mt) in 1950, the global annual production of plastic reached 465 Mt in 2019. Approximately 82 Mt is mismanaged and littered, resulting of 22 Mt plastic material that ends up in the environment.^1^ In nature, those plastics are degraded in micro- (MPL) and nano-particles (NPL) of plastic, exposed to UV rays, temperature variation, and oxidation resulting in weathered NPL. European Food and Safety Authority defines MPL and NPL as plastic particle between 0.1 and 5000 µm and 1 and 100 nm in size respectively.

Omnipresence of plastic in the environment and food products questioned about human exposure and hazardous property of MPL and NPL. This exposition mainly occurs through inhalation and ingestion.^2^ Indeed, MPL and NPL have been found in many dietary products like seafood, honey, sugar, salt, drink water^3^ and meat^4^ because of environmental or food containers contamination. These MPL and NPL enter in the human body through the ingestion of contaminated food, pass through the digestive tract and are released in faeces.^5^ Previous study have estimated annual human consumption of MPL at a ranges of 39000 to 52000 particles.^3^ However, NPL are not easily detectable and many pitfalls need to be avoid, like cross contamination.^6^ The physical and chemical properties of MPL and NPL, especially at nano size, facilitates their ability to cross intestinal epithelial barrier and previous studies found plastic particles in human tissues.^7–14^

So far, most of the studies for MPL and NPL detection have been performed on MPL identifying polyester (PES) as the main contaminant.^15^ However a recent paper analyzed MPL and NPL in water bottle by hyperspectral stimulated Raman scattering (SRS) imaging and showed that MPL and NPL were estimated to 10^5^ particles per liter with 90% at nanoscale mainly represented by Polystyrene (PS).^16^ In addition, an analysis of soil estimated a plastic contamination up to 100 mg/kg of soil and the most representative plastic was PS questioning its incorporation in food chain.^17^ Furthermore, extruded and expanded polystyrene have been widely used in food packaging due to their properties to maintain temperature and protect from oxygen. However, PS leakage to meat have been highlighted and question its consequence on costumers health.^18^

In parallel to increasing production and use of plastics, non-communicable diseases (NCD) are rising worldwide.^19^ Among NCD, adverse food reactions and Inflammatory Bowel Diseases (IBD) are associated with a rupture of tolerance of intestinal immune system toward food antigens and gut microbiota, respectively. NCD are multifactorial, involving genetic and environmental factors. Both disorders could find their origins in perinatal development based on the concept of Developmental Origin of Health and Diseases (DOHaD).^20–22^ Indeed, the perinatal period represents a highly dynamic transition time facilitating the maturation of the intestinal epithelium, the mucosal immune system and the intestinal microbiota and thus allowing the establishment of a homeostatic state throughout life. Presence of MPL in placenta and breast milk suggests that children are exposed to this pollutant during this critical developmental period.^14,23^

This study, based on mouse models aim to identify the consequences of perinatal exposure to pristine (PS50) or weathered polystyrene 50 nm (PS50w) on oral tolerance establishment and IBD severity.

## Materials and methods

### Particle preparation and characterization

Polystyrene particles, 50 nm, were obtained from Polysciences (Polybeads® 15913 -10 PS-COOH 50nm). They were weathered for 96 h in environmental conditions in a Q-SUN Xe-1 assay chamber (Q-Labs, Bolton, UK), using the weathering parameters indicated in the 4892-1 (2000) and 4892-2 (2013) ISO norms, i.e., an irradiance of 1.44 W.m-².nm-1 at 420 nm, a temperature of 40°C and a daylight filter. This corresponds to weather conditions of a sunny day at noon in equatorial region.

### Mice

Nulliparous female and male C3H/HeNRj (Janvier Labs, Le Genest St Isle, France) were housed in specific pathogen free (SPF) experimental at zootechny of US006/CREFRE, under 12h light-dark cycle, at constant temperature (23°C +/- 1°C). Animals were fed with SAFE® « A04 entretien » and watered with filtered tap water *ab libitum*, in cage with SAFE® « A04 entretien » litter. Pathogen-free conditions were monitored every 6 months in accordance with the full set of FELASA high standard recommendations.

### Study approval

All experimental procedures were approved by the French « Ministère de l’Enseignement Supérieur, de la Recherche et de l’Innovation » (project authorization number: APAFIS #34331-2021121309394360 v4) and were conducted in accordance with the European directive 2010/63/UE.

### Perinatal exposure to polystyrene 50nm

Animal experimental design is sum up in Figure 1. 3 females were bred with 2 males during 4 days, then males were removed. At 15th day of gestation (G15), gestational females were identified, individually isolated and randomly assigned to an experimental group. Gestational females were orally force fed every day from G15 to weaning (Post Natal Day PND 21) with 1.25mg of polystyrene 50nm (PS50) or 1.25mg of weathered polystyrene (PS50w) or vehicle (Mili-Q® purified water) (e.i., 0.35 mg/g of body weight (g bw)/week for a 25 g mice). Sex-ratio was monitored and litters size were standardized at 6 +/- 1 puppies per litter. Undercount or unisex litters were excluded from protocol and supernumerary offspring were sacrificed at PND1. At weaning (PND21), litters from the same experimental group of perinatal exposure are mixed and assigned to either inflammatory bowel diseases model or adverse food reaction model (oral tolerance or immunization). Mums were euthanized, jejunum, colon and sera were sampled.

**Figure 1.**
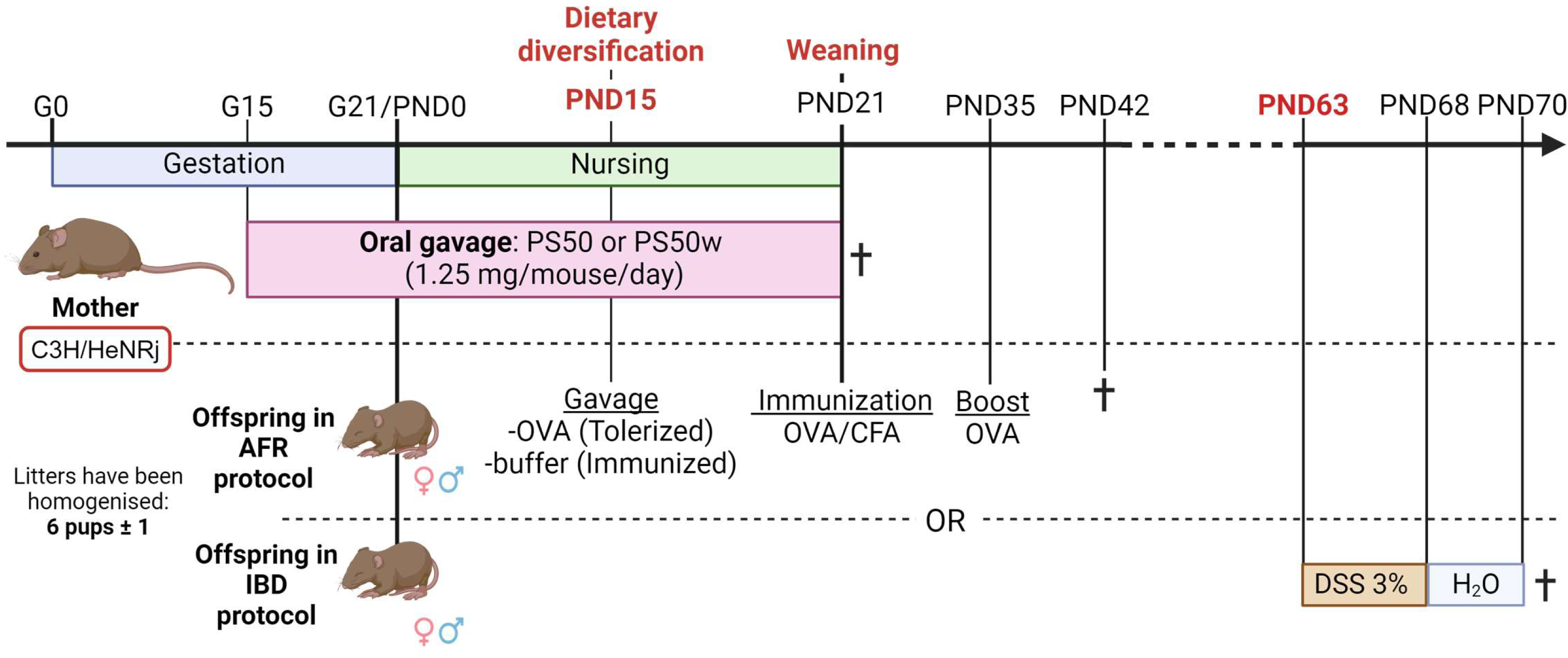
Experimental design of perinatal exposure to nano-particles of polystyrene. C3H/HeNRj mice were perinatally exposed to 50nm pristine polystyrene (PS50) or weathered polystyrene (PS50w) via their mother, that were daily force-fed from days 15^th^of gestation (G15) to weaning (post-natal day PND21) with 1.25 mg of PS50, PS50w or water. At birth litter were homogenized at 6 pups +/- 1. Thereafter, offspring were randomly included in adverse food reaction (AFR) or inflammatory bowel diseases (IBD) protocol. Mice in AFR protocol were force-fed with Ovalbumin (OVA) (tolerized group) or water (immunized group). Then offspring were immunized at PND21 with OVA in presence of Complete Freund Adjuvant (CFA) and immunized again (boost) at PND 35 before sacrifice one week later (PND42). At PND63, adult offspring in IBD protocol were exposed to 3% Dextran-Sodium Salt (DSS) in drinking water for 5 days, followed by 2 days of clearance with only drinking water to induce colitis before sacrifice.

### Model of Adverse food reaction (AFR)

At PND15, offspring perinatally exposed to PS50, PS50w, or vehicle (controls), were tested for oral tolerance induction (OVA-tolerized) and immunization (OVA-immunized) as indicated in Table 1. Briefly, OVA-tolerized pups were force-fed with 20 mg ovalbumin from chicken egg white Grade V (OVA) (Sigma-Aldrich®, A5503-10G) in 100µl of drink water at PND15 (Table 2) while OVA-immunized pups were orally given water alone. After 1 week (PND21), all pups were primed subcutaneously with 25 µg OVA in 1:1 complete Freund adjuvant (CFA): PBS and boosted 2 weeks later (PND35) by subcutaneous injection of 10 µg OVA. Sera and feces from offspring were collected and immediately stored at −80°C for biochemical analysis. Body weight of the offspring was measured.

**Table 1.** Summary of the treatments and doses used in the model of adverse food reaction. CFA: Complete Freund adjuvant; OVA : ovalbumin; sc: Subcutaneous; PND: Post Natal Day

| Age of mice | PND15 | PND21 | PND35 | PND42 |
| --- | --- | --- | --- | --- |
| OVA-tolerized | Gavage OVA 20mg<br>in drinking water | OVA in CFA<br>25µg sc | OVA boost<br>10µg sc | Sacrifice<br>Humoral response |
| OVA-immunized | Gavage drinking<br>water |  |  |  |

**Table 2.** Disease activity index. DAI: Disease activity index; max: maximum

| **Score DAI** | **% Weight loss** | **Consistency of feces** | **Blood in feces/anus** | **Mouse appearance** |
| --- | --- | --- | --- | --- |
| 0 | 0 | normal | no | normal |
| 1 | 0-5 | unformed | yes | bristly |
| 2 | 5-10 | light diarrhea |  | prostrate |
| 3 | 10-15 | heavy diarrhea |  | prostrate and<br>bristly |
| 4 | 15-20 |  |  | lethargic |
| max = 12 | 4 | 3 | 1 | 4 |

### Model of colitis

To test susceptibility to DSS-induced colitis of 63-days old mice perinatally exposed to PS50 or PS50w offspring were exposed to 3% (w/v) Dextran Sodium Salt (DSS) (MP Biomedical®, 160110) in their drinking water for 5 days, followed by 2 days of clearance with tap water. Control mice received drinking water without DSS. Colitis severity was scored by daily observation of the Disease Activity Index (DAI), composed of the following parameters (Table 2): weight loss, stool consistency, blood in stool and, mice appearance with a maximum score of 12, adapted from the scoring of Cooper et al.^24,25^ After 7 days, offsprings’ feces were collected then stored at −80°C, and were euthanized to collect ileum and colon which were immediately frozen in liquid nitrogen and stored at −80°C for future biochemical analysis. At sacrifice, macroscopic observation of intestinal inflammation was measured using the following parameters (Table 3): colon length, colon thickness, oedema size, presence of hyperemia and mesenteric adherences; presence of stool in the colon, with a maximum score of 17, adapted from the scoring of Wallace et al.^26^

**Table 3.** Macroscopic damage score. max: maximum

| Macroscopic score | Colon length (cm) | Colon thickness (mm) | Oedema size | Hyperemia | Adherence | Feces in colon |
| --- | --- | --- | --- | --- | --- | --- |
| 0 | $x \geq 7.5$ | $x < 0.3$ | 0 | absence | absence | yes |
| 1 | $7.5 > x \geq 6.5$ | $0.35 > x \geq 0.3$ | $0.5 \geq x > 0$ | presence | low | No |
| 2 | $6.5 > x \geq 6$ | $0.40 > x \geq 0.35$ | $1 \geq x > 0.5$ | - | medium | |
| 3 | $6 > x \geq 5.5$ | $0.45 > x \geq 0.40$ | $1.5 \geq x > 1$ | - | high | |
| 4 | $x < 5.5$ | $x > 0.45$ | $x > 1.5$ | - | - | |
| max = 17 | 4 | 4 | 4 | 1 | 3 |  |

### Measurements of colonic permeability

*Ex vivo* permeability assays were done in Ussing chamber systems as previously described.^27^ Colonic fragments of offspring in the colitis protocol were mounted in Ussing chambers (Physiologic Instruments, San Diego, CA, USA). Tissues were bathed 1h with oxygenated thermostated Ringer’s solution. Ringer’s solution is composed of 6.72g of NaCl, 2.1g of NaHCO3, 50 ml of solution with 4.82 g of MgCl2 (6H2O) and 3.52g of CaCl2 (2H2O), 100 ml of solution with 4.16g of K2HPO4 and 0.54g of KH2PO4, qsp 1 L of distilled water. Fluorescein isothiocyanate–dextran (FITC-4kDa) 0.4 mg/ml (FD4; Sigma-Aldrich®) was added to mucosal compartment. Epithelial permeability to FITC-4kDa was determined by measuring the fluorescence intensity (FI) 485 nm/525 nm in the serological compartment, using an automatic EnSight™, Multimode microplate reader. Permeability was calculated as the ratio of flux/concentration, and expressed as pmol/h/cm^2^ as previously described.^28,29^

### Feces supernatant

Feces were lysed in 1 ml of PBS with proteases inhibitor using Lysing Matrix D tubes (fastprep ref: 6913500) and Precellys 24 Touch homogenizer. The homogenate was centrifuged at 4°C for 10 minutes at 1600 g. Protein concentrations was measured in supernatant with BCA uptima kit (Interchim, Montlucon, France).

### Serum Preparation Protocol

At sacrifice, blood was collected intracardially without heparin, centrifuged for 10 minutes at 1000 g and serum collected and kept at −80°C.

### Humoral response in feces and plasma

Plates maxisorp were coated with 5 µg/ml of sheep anti-mouse IgA (Sigma) or goat anti-mouse IgG (SouthernBiotech, Cliniscience, Nanterre, France), incubated plasma or feces supernatant, detected with 1.5 µg/ml HRP-conjugated goat anti-mouse IgA (Sigma) or goat anti-mouse IgG (SouthernBiotech), HRP was revealed using TMB and the reaction was stopped with H_2_SO_4_ before reading at 450 nm using automatic Varioskan™ microplate reader.

### ELISA Titration Protocol for Anti-Ovalbumin IgG/IgG1/IgG2a Detection

The ELISA titration protocol was performed to quantify anti-ovalbumin Ig. Ovalbumin antigen was coated onto a 96-well Nunc MaxiSorp™ plate, followed by blocking with 5% SVF in PBS. Serial dilution of serum samples were then added to the wells and incubated. HRP-conjugated anti-mouse IgG/IgG1/IgG2a (SouthernBiotech, Cliniscience, Nanterre, France) detection antibody was applied. TMB substrate was added to develop the colorimetric reaction. The reaction was stopped with H_2_SO_4_, and the absorbance was measured at 450 nm using a Varioskan™.

### Calprotectin measurement in feces

The measurement of calprotectin was determined in feces supernatant with a Duoset ELISA kits (S100 Calcium Binding Protein A8 - S100A8).

### Statistical Analysis

Statistical analyses were performed using GraphPad Prism 10.00 (GraphPad software, San Diego, CA) software package for PC. Results are summarized in box-and-whiskers plots. The down edge of the box represents the lower quartile, the up edge the upper quartile and the splitting line of the box the median. The upper whiskers represent the maximum value whereas the lower whiskers represent the minimum value. All kinetic charts are expressed as mean ± SEM. Multigroup comparisons were performed using a One-way analysis of variance. Two-group comparisons were performed using 2-way analysis of variance. A value of P <0.05 was considered statistically significant. All P values were 2-sided.

## Results

### Physico-chemical characterization of the PS particles

Transmission electron microscopy images of pristine and weathered particles are shown in Figure 2A-B respectively. Their size distribution is reported in Figure 2C-D. Pristine and weathered particles were characterized in terms of their average hydrodynamic diameter and zeta potential, as reported in Figure 2E. Their mean hydrodynamic diameter were 49.96±0.61 (polydispersity index 0.053±0.010) and 49.9±1.29 nm (polydispersity index 0.048±0.004) for pristine and weathered particles, respectively, and their zeta potential −34.2±2.9 mV and −30.0±1.2 mV, respectively.

**Figure 2.**
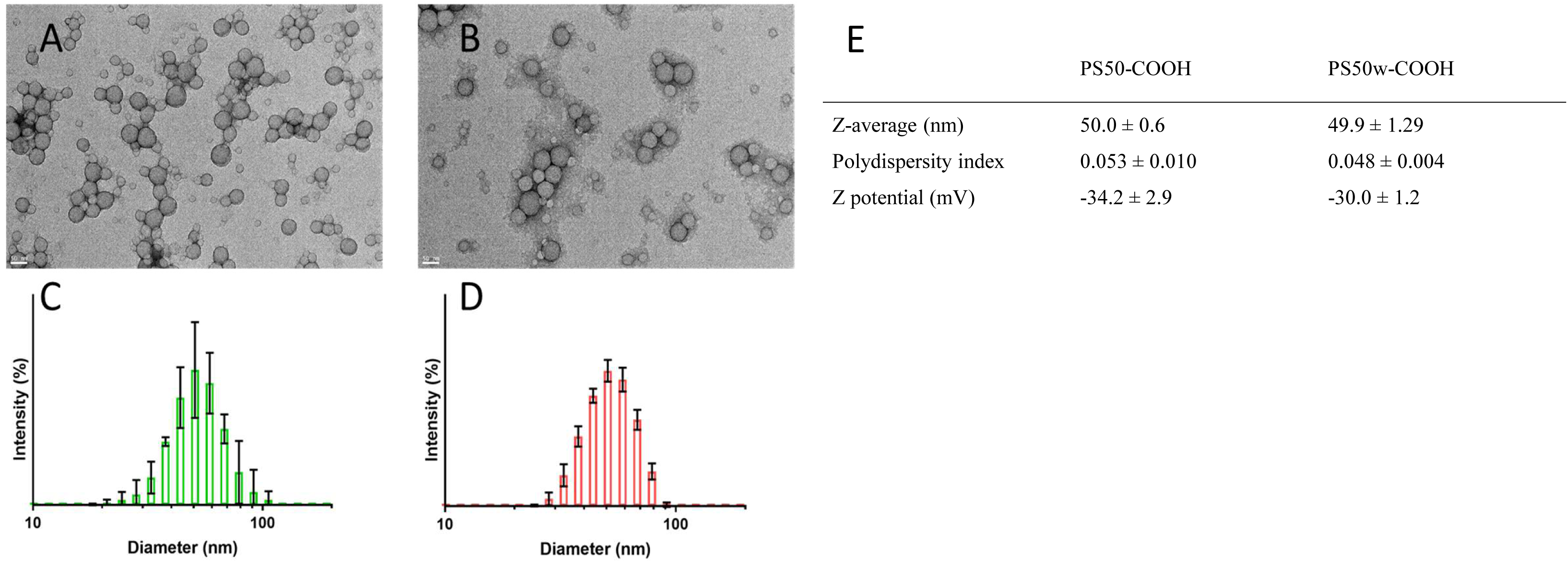
Shape and size of PS particles. Electronic microscopy images (A, B) and size distribution as measured via dynamic light scattering (C, D) of pristine PS50-COOH nanoparticles (A, C) or weathered PS50-COOH nanoparticles (B, D).

### At weaning, female offspring weight is increased by perinatal exposure to PS50w

During the week of gestational oral exposure to pristine (PS50) or weathered (PS50w) nanoparticles of Polystyrene (PS), a tendency to higher gestational mortality was observed in females orally exposed to PS50 even thought this effect is not statistically significant (Figure 3B). At birth, the number of offspring per litter and the sex ratio is not affected by oral perinatal exposure to PS50 nor PS50w (Figure 3C). At weaning (Post Natal Day; PND21), females offspring weight was increased by perinatal exposure to PS50w (Figure 3D). However, at weaning (lactating day 21; L21), after 4 weeks of daily oral gavage with PS50 or PS50w, mothers’ weight was not affected nor intestinal length nor IgG and IgA humoral response in feces (Figures 3E-I). In addition, serosal IgG concentration were not modified by 4 weeks daily oral gavage with PS50nm nor PS50w (Supplementary Figure 1B). Finally, blood glucose concentration at fasted and fed status was not altered by 4 weeks daily oral gavage with PS50 nor PS50w (Supplementary Figure 1C).

**Figure 3.**
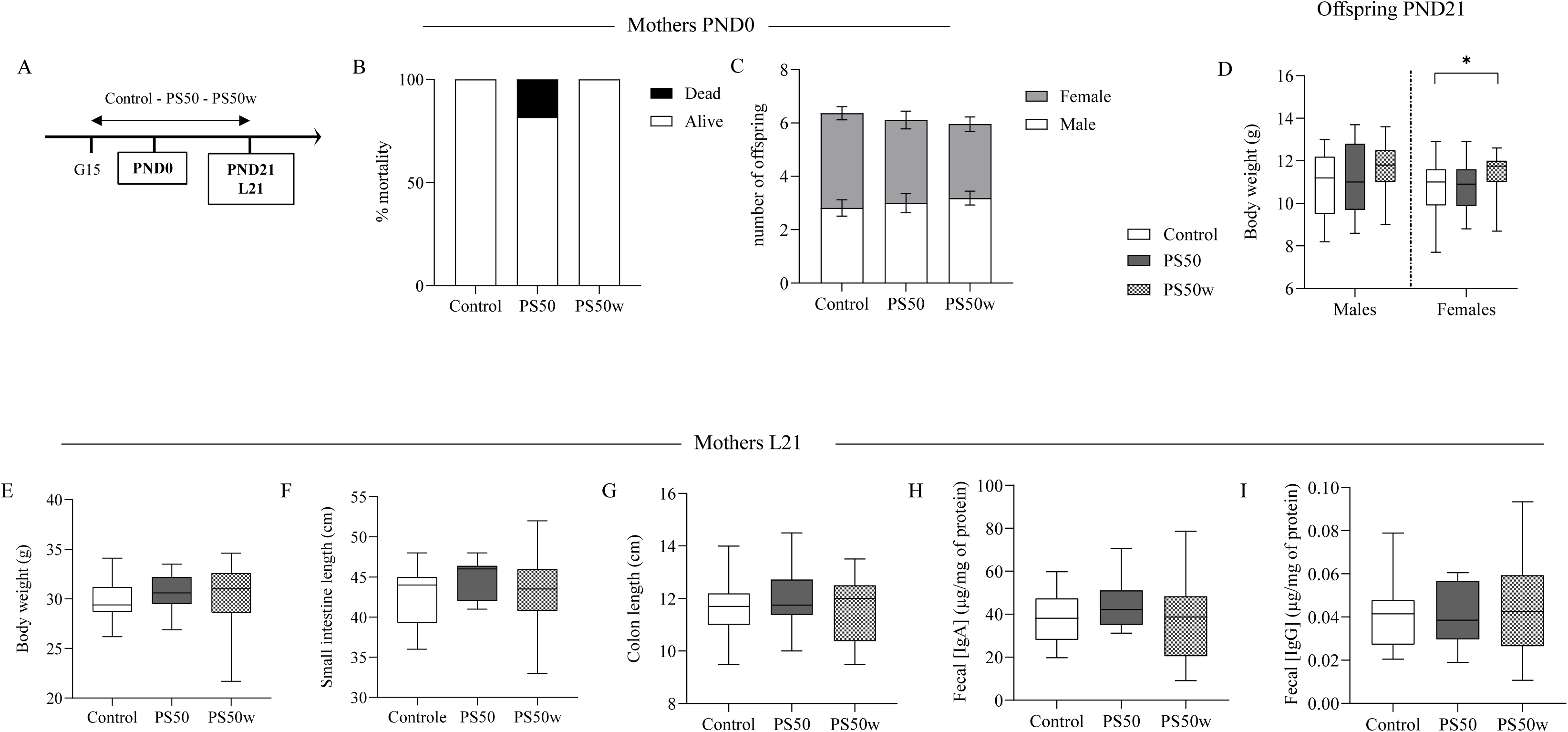
At weaning females offspring weight is increased by perinatal exposure to PS50w. (A) Mice were perinatally exposed to 50nm polystyrene (PS50) or weathered polystyrene (PS50w) via their mother, that were daily force-fed from days 15^th^of gestation (G15) to weaning (post-natal day 21, PND21lactating day 21, L21) with 1.25 mg of PS50, PS50w or water. (A-I) Several parameters have been determined at postnatal day 0 (PND0, B and C) in mothers, and postnatal day 21 (PND21) in offspring (D) and lactating day 21 in mothers (E-I). The following parameters have measured: (B) Percentage of female mice that died during the gestational period, (C) number of pups per litter and the sex ratio, (D) body weight of offspring, (E) body weight of mothers, (F) small intestine’s length of mothers, (G) large intestine’s lenght of mothers, (H) mother’s fecal concentration of (H) IgA and (I) IgG. n=10 mice minimum per group, for offspring at least 2 independent litters were analyzed.

### Oral tolerance establishment in offspring at postnatal day 15 is not affected by perinatal exposure to PS50 nor PS50w

First, we analyzed the consequences of daily oral gavage with PS50 or PS50w on mothers and offspring at L15/PND15 (Figures 4B-D). IgA nor IgG fecal concentrations were affected by daily oral gavage with PS50 nor PS50w in mothers at L15 (Figures 4B and C). However, perinatal exposure to PS50w but not PS50 increased fecal IgA in male and female offspring at PND15 (Figure 4D).

**Figure 4.**
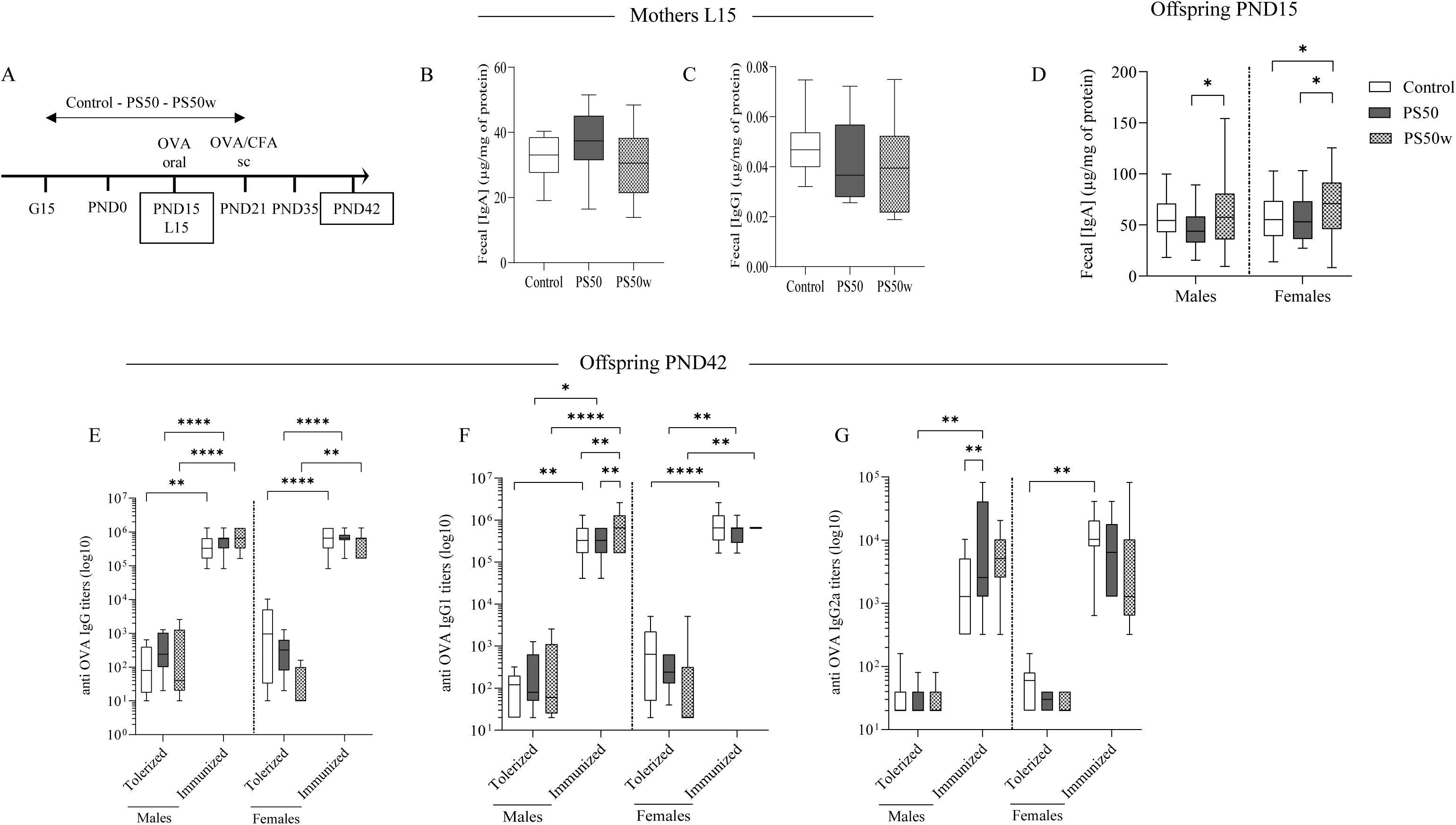
Oral tolerance establishment in offspring at postnatal day 15 is not affected by perinatal exposure to PS50 nor PS50w. (A) Mice were perinatally exposed to 50nm polystyrene (PS50) or weathered polystyrene (PS50w) via their mother, that were daily force-fed from days 15^th^of gestation (G15) to weaning (post-natal day 21; PND21) with 1.25 mg of PS50, PS50w or water. At lactating day 15 (L15), the fecal concentration of (B) IgA and (C) IgG from mothers have been monitored. (D) At PND15, the fecal concentration of IgA from offspring have been measured. (E) Then, at PND42, (E) anti-OVA IgG, (F) anti-OVA IgG1 and (G) anti-OVA IgG2 titers were measured in serum. *p<0.05; **p<0.01; ***p<0.001 and ****p<0.0001 vs. indicated group; n=10 mice minimum per group, for offspring at least 2 independent litters were analyzed

We then investigated if perinatal exposure to PS50 or PS50w could impair oral tolerance establishment and immune response to immunization at solid food introduction and vaccination time frame (Figures 4E-G). Oral tolerance is characterized by the absence of humoral response (anti-OVA IgG) to the orally administered antigen (OVA). In control offspring (i.e., not exposed to PS50 nor PS50w), OVA tolerization (i.e., OVA oral ingestion prior to OVA systemic immunization) significantly decreased by 1 000-fold the serum anti-OVA IgG titers, validating our protocol for induction of oral tolerance to OVA at PND15 for the first time (Fig. 4E). Oral tolerance establishment tested at PND15 was not affected by PS50 nor PS50w perinatal exposure in male and female offspring. Immune response to immunization (mimicking vaccination) was not affected by PS50 nor PS50w perinatal exposure in male and female offspring (Figure 4E). We then tried to decipher the response between Th2 (IgG1 - Figure 4F) and Th1 (IgG2a - Figure 4G). In immunized male offspring, perinatal exposure to PS50w increased anti OVA IgG1 titers compared to control and PS50 group (Figure 4F) whereas perinatal exposure to PS50 increased anti OVA IgG2a titers compared to control group (Figure 4G).

### DSS-induced colitis is exacerbated by both perinatal exposure to PS50 and PS50w with stronger effect of PS50w in male offspring

First, we analyzed the long-lasting consequences of perinatal exposure to PS50 or PS50w on male offspring at postnatal day 56 (PND56), one week before colitis induction. Perinatal exposure to PS50 nor PS50w impacted fasted (Supplementary Figure 2B) or fed glycemia (Supplementary Figure 2C) of male offspring. However, weight of male offspring was increased by perinatal exposure to PS50w but not PS50 (Supplementary Figure 2D). Then, we investigated if perinatal exposure to PS50 or PS50w could aggravate DSS-induced colitis at PND63. Increase of body weight in PS50w was still observed at PND63 (beginning of DSS-induced colitis protocol) (Figure 5B). In male offspring, DSS increased the weight loss (Figure 5C) and the Disease Activity Index (DAI) (Figure 5D), and increase the score of mice appearance (Supplementary Figure 3B) and the feces consistency score (Supplementary Figure 3C), without any modification by perinatal PS50 nor PS50w perinatal exposure. DSS-induced colitis increased fecal IgA (Supplementary Figure 4B), IgG (Supplementary Figure 4C) and calprotectin (Supplementary Figure 4E) and induced intestinal hyper-permeability to FD4 (Supplementary Figure 4D) without alteration by perinatal exposure to PS50 nor PS50w. However, perinatal exposure to PS50 increased DSS-induced macroscopic score of colitis compared to control group and PS50w increased DSS-induced macroscopic score of colitis compared to control and PS50 group (Figure 5E). Among the factors used to calculated macroscopic score, increase of edema size seems to be the main driver of the effect (Figure 5H). Looking in detail, perinatal exposure to PS50 increased DSS-induced adherence score compared to control group (Figure 5I). Perinatal exposure to PS50w increased DSS-induced adherence score (Figure 5I) and colon thickness (Figure 5G) and decrease colon length (Figure 5F) compared to control group. Perinatal exposure to PS50w also increased DSS-induced edema size (Figure 5H) and colon thickness (Figure 5G) compared to PS50 group.

**Figure 5.**
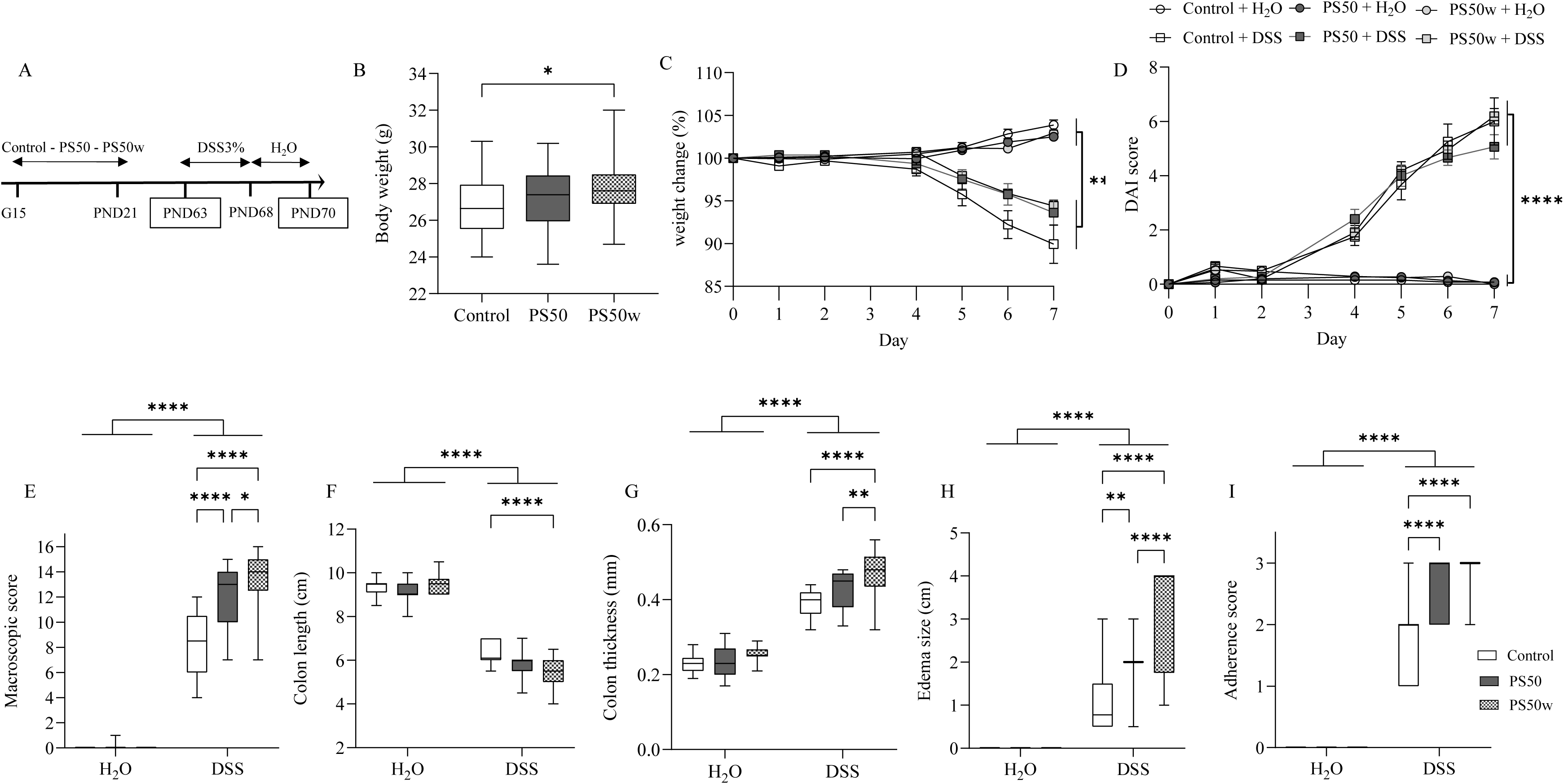
DSS-induced colitis is exacerbated by both perinatal PS50 and PS50w exposure with stronger effect by PS50w in males offspring. (A) Mice were perinatally exposed to 50nm polystyrene (PS50) or weathered polystyrene (PS50w) via their mother, that were daily force-fed from days 15^th^of gestation (G15) to weaning (post-natal day 21) with 1.25 mg of PS50, PS50w or water (A) At PND63, mice have been (B) weighed and colitis has been orally induced by introducing Dextran Sulfate Sodium (DSS) into drinking water at 3% for 5 days followed by 2 days of regular water. During the experimental DSS procedure (7 days), (C) body weight and the (D) disease activity index (DAI) have been monitored daily. (E-I) Then, at the end of the DSS procedure (PND70), mice have been sacrificed and (E) macroscopic score, (F) colon length and (G) thickness, (H) edema size and (I) adherence score have been monitored. n=10 mice minimum per group from at least 2 independent litters.

### DSS-induced colitis is exacerbated by both perinatal exposure to PS50 and PS50w in female offspring

First, we analyzed the long-lasting consequences of perinatal exposure to PS50 or PS50w on female offspring at PND56 (one week before colitis induction). Fasted (Supplementary Figure 2B) and fed glycemia (Supplementary Figure 2C) were not modified by PS50 nor PS50w perinatal exposure. However, weight of female offspring was increased by perinatal exposure to PS50w at PND56 (Supplementary Figure 2D) compared to control and PS50 groups. Then, we studied if perinatal exposure to PS50 or PS50w could aggravate DSS-induced colitis at PND63. Increase of body weight in PS50w was still observed at PND63 (beginning of DSS-induced colitis protocol) (Figure 6B). DSS induced weight loss (Figure 6C) and increased the Disease Activity Index (DAI, Figure 6D). Perinatal exposure to PS50w increased DAI at day 5 compared to control and PS50 groups (Figure 6D). Among the actors of the DAI both mice appearance (Supplementary figure 3D) and feces consistency (Supplementary figure 3E) were significantly modified suggesting a multifactorial cumulative effect. DSS-induced colitis increased fecal IgA (Supplementary Figure 4F), and IgG (Supplementary Figure 4G), induced intestinal hypermeability to FD4 (Supplementary Figure 4I) without alteration by perinatal exposure to PS50 nor PS50w. However intestinal permeability is decreased by PS50w perinatal exposure in DSS-induced colitis group compared to control DSS and PS50-DSS. DSS increased fecal calprotectin concentration which is enhanced by PS50 perinatal exposure (Supplementary Figure 4H). However, perinatal exposure to PS50 and PS50w increased DSS-induced macroscopic score of colitis compared to control group (Figure 6E). Among the factors used to calculated macroscopic score increase of edema size (Figure 6H), mesenteric adherences (Figure 6I) and colon thickness (Figure 6G) and decrease of colon length (Figure 6F) appeared to be the main driver of the effect.

**Figure 6.**
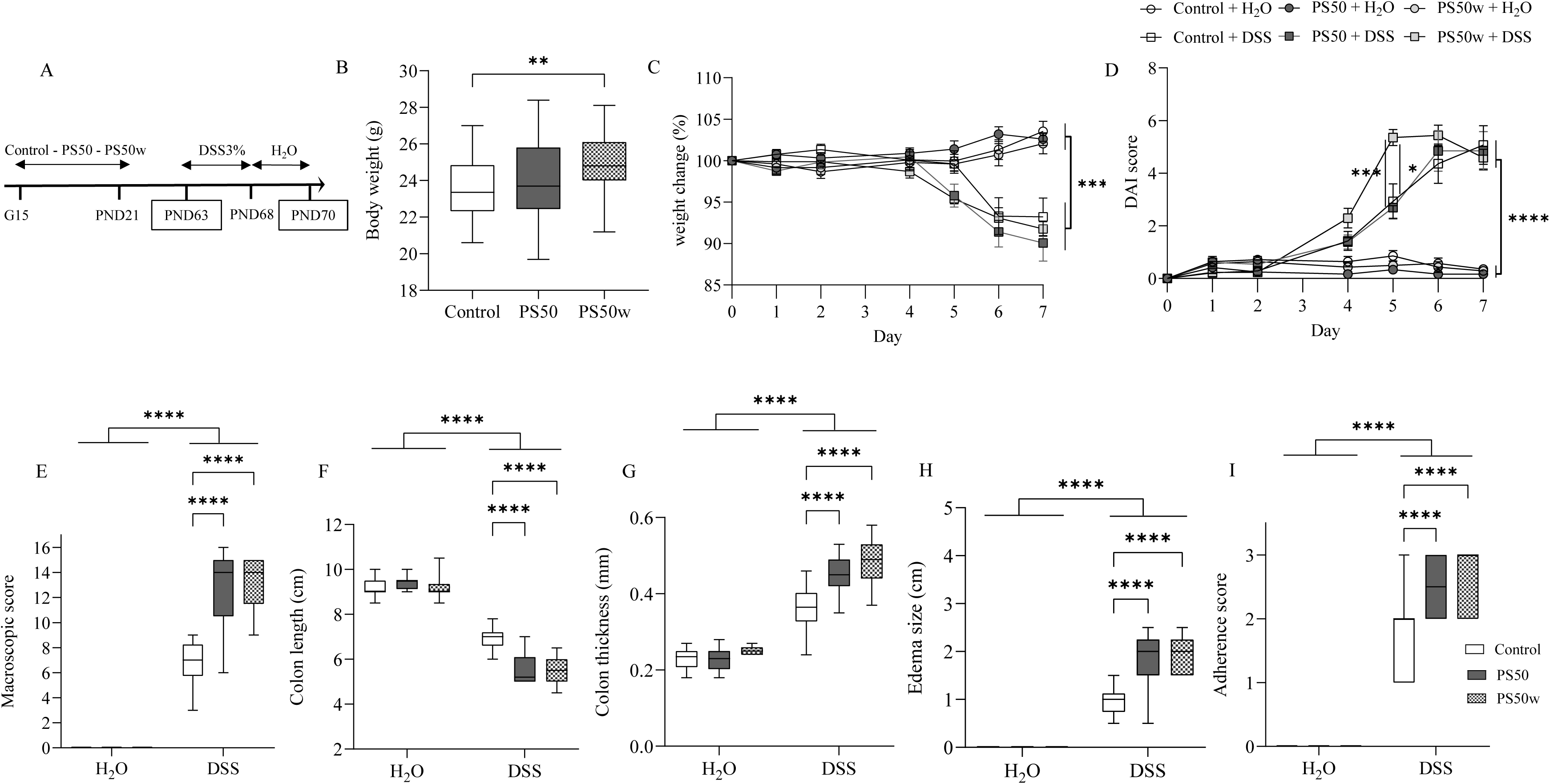
DSS-induced colitis is exacerbated by both perinatal PS50 and PS50w perinatal exposure in female offspring. (A) Mice were perinatally exposed to 50nm polystyrene (PS50) or weathered polystyrene (PS50w) via their mother, that were daily force-fed from days 15^th^of gestation (G15) to weaning (post-natal day 21; PND21) with 1.25 mg of PS50, PS50w or water. (A) At PND63, mice have been (B) weighed and colitis has been orally induced by introducing Dextran Sulfate Sodium (DSS) into drinking water at 3% for 5 days followed by 2 days of regular water. During the experimental DSS procedure (7 days), (C) body weight and the (D) disease activity index (DAI) have been monitored daily. (E-I) Then, at the end of the DSS procedure PND70, mice have been sacrificed and (E) macroscopic score, (F) colon length and (G) thickness, (H) edema size and (I) adherence score have been monitored. n=10 mice minimum per group from at least 2 independent litters.

## Discussion

Given the widespread distribution of nanoplastics in our environment and human food chain, there is an urgent need to study their potential health outcomes especially on non-communicable diseases onset. Perinatal exposure through gestation and lactation might represents a critical window corresponding to foetus and neonate maturation. In this original work we addressed 2 main questions: i) did perinatal exposure to pristine or weathered nano- particles of polystyrene impair oral tolerance establishment of oral tolerance and immune response at solid food introduction, ii) did perinatal exposure to pristine or weathered nano- particles of polystyrene had long lasting consequences on IBD onset on adulthood? At steady state, we showed that pristine nor weathered polystyrene nanoparticles impact intestinal, metabolic and immunological parameters of gestational and lactating mice. However, as speculated, foetus and neonates indirectly exposed to PS50 or PS50w had long lasting modification of numerous physiological parameters observed in male and female. Indeed, all along the life course, perinatal exposure to PS50 and/or PS50w has consequences on body weight, faecal IgA concentration on basic conditions and on colitis induced experimental model perinatal exposure to nano-polystyrene pristine and weathered worsen colitis and intestinal hyperpermeability triggered by colitis. This study highlights a sexual dimorphism as the observed effects were more pronounced in females than in males.

Compare to MPL, the small size of NPL makes them difficult to quantify and identify in the environment^30^ which means they represent an invisible threat for environmental, animal and human health. In our work we wanted to compare the consequences of perinatal exposure of pristine and weathered nano-particles of PS50 in order to mimic the degradation of plastics during its transfer in the environment and explore the potential toxic effect. Gestating and lactating mothers were exposed via oral route as food^31^ and beverage^3^ represent a main sources of exposure. Current human exposure of MPL is estimate to 5 g/week for adults which is represent approximately 0.07 mg/bw g/week, even though this value is still under debate and seem to have been overestimated.^32^ No data is available about the human exposure of NPL. Thus, based on the estimated human plastic exposure (MPL) and on the several doses of PS’s NPL exposure reported in the literature, we choose to expose mice at 1.25 mg of PS50 pristine or weathered per dam mice per day.

First, we analysed the consequences of perinatal exposure to PS50 or PS50w on mothers and offspring on basal conditions. During gestation perinatal exposure to PS50 or PS50w did not induce mortality. At birth perinatal exposure to PS50 or PS50w did not affect sex ratio. At PND15, solid food introduction before AFR protocol perinatal exposure to PS50w increased fecal IgA concentration. These results suggest that indirect exposure to PS50w via dam mice have induced a higher intestinal immune response in male and female offspring. At weaning (PND21), perinatal exposure to PS50w increased body weight of female offspring. No modification of metabolism as body weight or fed and fasted glycemia were observed in force fed mothers. However, it has been reported that mice exposed to NPL of PS presented an increased plasmatic glucose level with no longer modification of plasma insulin, then suggesting that hyperglycaemia was linked to insulin resistance. However, these alterations were induced by one administration of very high dose of PS corresponding to the annual exposure of human in one single dose.^33,34^ At PND56 and PND63 our data have evidenced that perinatal exposure to PS50w increased the body weight in male and female offspring. Those data agree with the study of Jeong et al. showing that early life exposure to NPL of PS increased the body weight in offspring.^35^ Still we could observe that body weight of females was increased as early as PND21 by PS50w. This sexual dimorphism had been already highlighted by the study of Jeong et al. reporting a different weight gain between male and female exposed to NPL’s PS during early life^35^ as well as on the secretion of sexual hormones including testosterone, luteinizing hormones, follicle-stimulating hormones.^36^

Then we wondered if perinatal exposure could affect oral tolerance establishment and immune response to vaccination at food diversification. No differences in the establishment of the oral tolerance was observed between control, PS50 or PS50w group. In the same way, PS50 nor PS50w affected immune response to vaccination. Nonetheless, one may notice that perinatal exposure to PS50w increased anti OVA IgG1 titers in male offspring whereas PS50 increased anti-OVA IgG2a in immunized groups suggesting higher immune responses.

Finally, we wondered if perinatal exposure to PS50 or PS50w had long lasting consequences on IBD severity. Concerning the impact of NPL’s and the intestinal homoeostasis, our data showed that only perinatal exposure of PS50 in female increased fecal calprotectin in DSS-induced colitis conditions. This increase is associated with higher colonic permeability. We did not observe any long-lasting consequences of NPL perinatal exposure on intestinal barrier at steady state.

Our data are supported by those of Schwarzfischer et al., indicating that exposure to PS NPL through drinking water (6 months at 0.056 mg/bw g/week) did not modify the colonic expression of cytokines and tight junctions.^37^ However, Lian and al reported that 28-days oral exposure with PS50 particles with a daily dose very closed to that used in our study, increased the ROS production by intestinal epithelial cell. Moreover, exposure of PS50 at elevated dose is reported to increase the number of apoptotic cells as well as the intestinal permeability.^38^

These discrepancies are probably due to the dose used, the duration of exposure and age of exposure.

Our data evidenced that perinatal exposure to PS50 or PS50w aggravated the colitis severity induced by DSS at adulthood. Perinatal exposure to PS50w induced a more deleterious impact on the colitis severity than PS50 in both males and females. Those data are in accordance with previous study reported a positive correlation between the concentration of MPL in stool and the severity of IBD.^39^ In inflammatory context, it has recently been shown that exposure to NPL’s PS (0.056 mg/bw g/week for a 25 g mice) in drinking water during 6 month did not aggravated the DSS induced colitis as well cytokines and tight junction expressions.^37^ This discrepancy may be due to the fact that this study was performed in adult and early life is a critical period for the establishment of intestinal homeostasis according to the DOHaD concept. Adult exposure to NPL’s PS (0.056 mg/bw g/week for a 25 g mice) did not modify the colitis severity induced by DSS,^37^ strengthening our hypothesis that early life exposure to NPL’s PS, through mother exposition, could trigger a deleterious imprinting of the gastrointestinal tract persisting throughout life.

Together, these data shown that early exposure to PS50 has a deleterious impact on the gut development during the perinatal period then altering intestinal homeostasis throughout life with a worsen effect by weathered PS50w.

## Conclusion

Our study demonstrates that early life exposure to nano-particles of polystyrene induces long-lasting consequences in intestinal homeostasis including enhanced susceptibility to develop colitis throughout life in male and female offspring. Moreover, our data also evidence that perinatal exposure to weathered nanoparticles of polystyrene triggers more deleterious on the gastrointestinal tract than the pristine nanoparticles of polystyrene. Finally, our results also show that the severity of digestive damage induced by perinatal exposure to nanoparticles differs according to sex, with much more severe effects in females than in males. These results emphasized the importance to investigate both sexes, and the importance of the plastic structure, since weathered PS have more deleterious impact on the intestinal health than virgin PS.

In conclusion our results provide new insights into the potential hazards of nanoparticles of polystyrene on non-communicable diseases and the consequences of maternal exposure to the nanoparticles of polystyrene on the health of the offspring by modifying the perinatal imprinting.

## Authors contributions

Study concept and design : MC, FB and SM; acquisition of data : MA, AR, LI, MC, FB and SM; analysis and interpretation of data: MA, AR, MC, EM, FB and SM; drafting of the manuscript: MA, AR, MC, FB and SM; critical revision of the manuscript for important intellectual content: MC, FB and SM; statistical analysis: MA, AR, FB and SM; obtained funding : MC, FB and SM; study supervision: FB and SM.

## Acknowledgements

We thank the ANINFIMIP EquipEx facility headed by Professor E Oswald, and supported by the French government through the Investments for the future programme (ANR-11-EQPX- 0003).

## Supplementary Figures legends

**Supplementary Figure 1. Perinatal exposure of PS50 or PS50w did not alter the quantity of IgG and the glycemia of mums.** (A) Mice were perinatally exposed to 50nm polystyrene (PS50) or weathered polystyrene (PS50w) via their mother, that were daily force-fed from days 15^th^ of gestation (G15) to weaning (Postnatal Day 21, PND21/L21) with 1.25 mg of PS50, PS50w or water. (A) at L21, the concentration of (B) IgG in the mother’s serum and the (C) glycemia have been determined. At least n=10 mice per group.

**Supplementary Figure 2. Perinatal exposure to PS50w increased body weight of offspring at PND56.** (A) Mice were perinatally exposed to 50nm polystyrene (PS50) or weathered polystyrene (PS50w) via their mother, that were daily force-fed from days 15^th^ of gestation (G15) to weaning (Postnatal Day 21, PND21) with 1.25 mg of PS50, PS50w or water. At post- natal day 56, (B) fasted and (C) fed glycemia as well as (D) the body weight have been monitored in both males and females offspring. n=10 mice minim per group, at least 2 independent litters.

**Supplementary Figure 3. Perinatal exposure to PS50w increased mice appearance score and feces consistency at day 5 of DSS-induced colitis.** (A) Mice were perinatally exposed to 50nm polystyrene (PS50) or weathered polystyrene (PS50w) via their mother, that were daily force-fed from days 15^th^ of gestation (G15) to weaning (Postnatal Day 21, PND21) with 1.25 mg of PS50, PS50w or water. (A) At PND63, colitis has been orally induced by introducing Dextran Sulfate Sodium (DSS) into drinking water at 3% for 5 days followed by 2 days of regular water. During the experimental DSS procedure (7 days), (B and D) the mice’s appearance as well as the (C and E) feces consistency have been daily monitored in males and female offspring. n=10 mice minim per group, at least 2 independent litters.

**Supplementary Figure 4. Perinatal exposure to PS50 or PS50w increased the susceptibility of male’s offspring to develop colitis later in life.** (A) Mice were perinatally exposed to 50nm polystyrene (PS50) or weathered polystyrene (PS50w) via their mother, that were daily force-fed from days 15^th^ of gestation (G15) to weaning with 1.25 mg of PS50, PS50w or water. (A) At PND63, colitis has been orally induced by introducing Dextran Sulfate Sodium (DSS) into drinking water at 3% for 5 days followed by 2 days of regular water. Then, at the end of the DSS procedure, mice have been sacrificed and fecal concentration of (B and F) IgA, (C and G) IgG and (D and H) of calprotectin have been studied in (B-D) in males and in (F-H) in female offspring. (E and I) The colonic permeability to dextran-FITC 4kDa (FD4) has been measured in Ussing chamber. n=10 mice minim per group, at least 2 independent litters.

