## Supplementary figures and images for "Deleterious imprinting of perinatal exposure to nano-polystyrene on inflammatory bowel diseases"

### Supplemental Figures

Supplementary figure 1

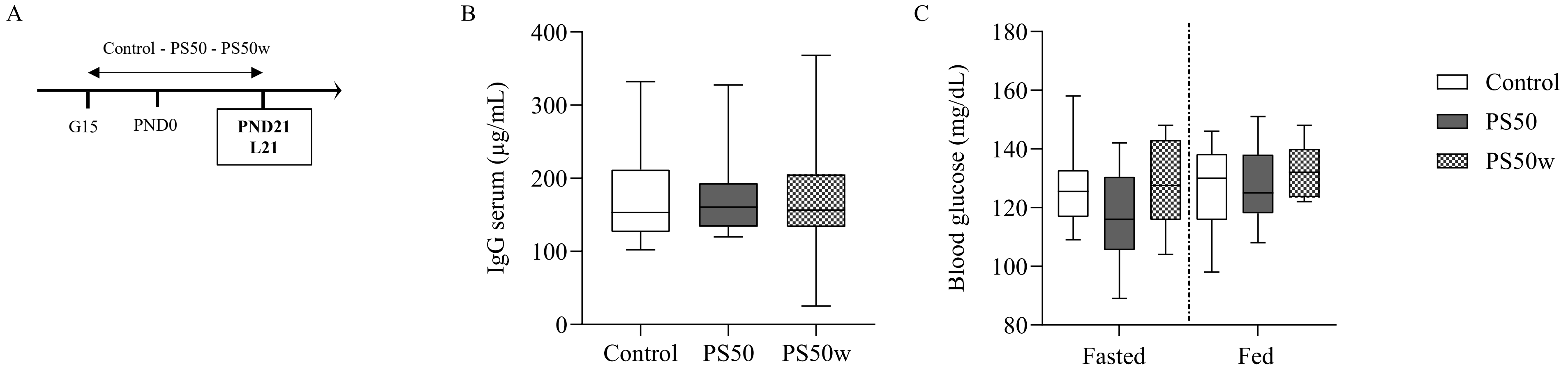

Supplementary figure 2

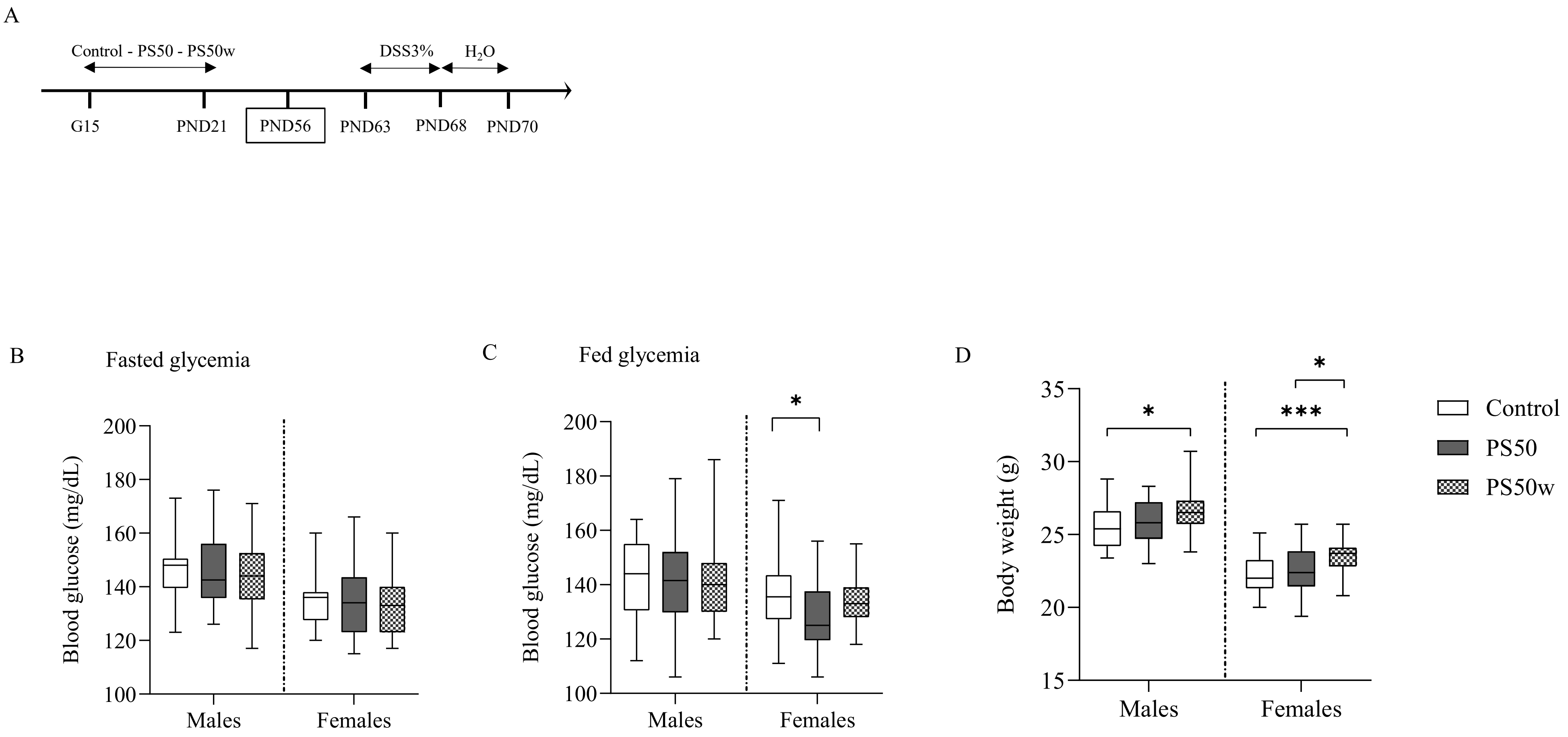

Supplementary figure 3

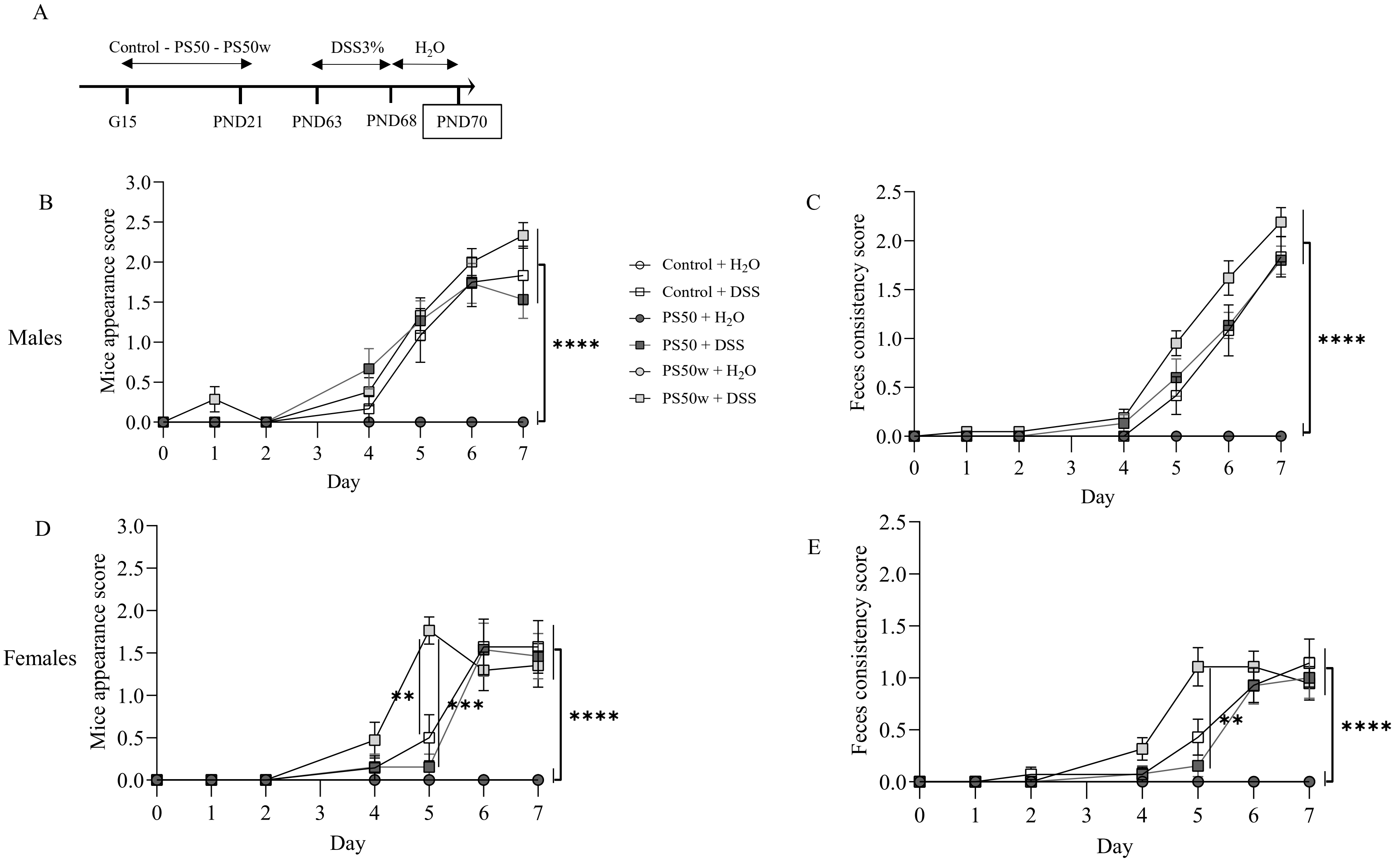

Supplementary figure 4

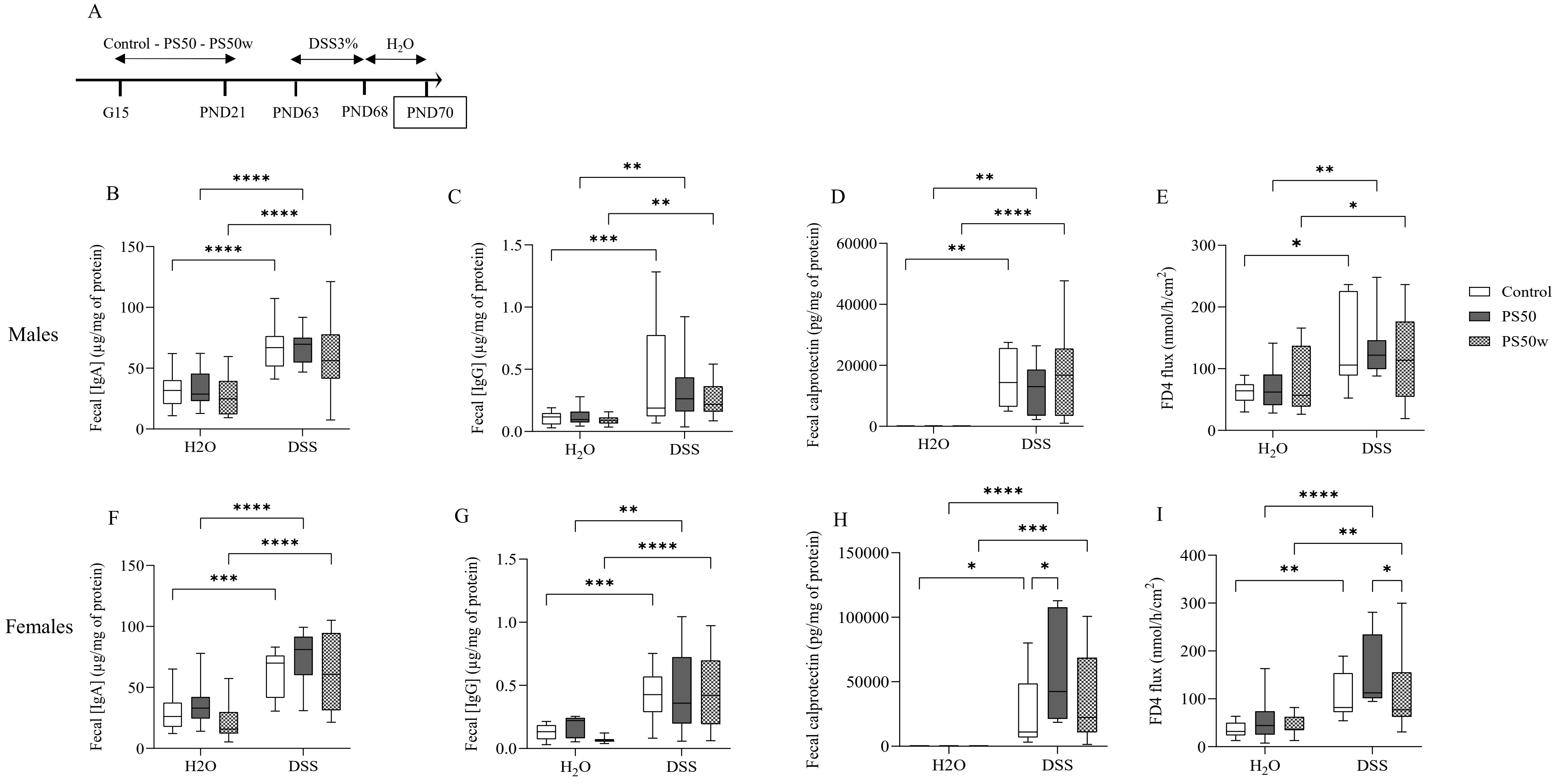
